# Phase variation-based biosensors for bacteriophage detection and phage receptor discrimination

**DOI:** 10.1101/851808

**Authors:** David R. Olivenza, Josep Casadesús, Mireille Ansaldi

## Abstract

Environmental monitoring of bacteria using phage-based biosensors has been widely developed for many different species. However, there are only a few available methods to detect specific bacteriophages in raw environmental samples. In this work, we developed a simple and efficient assay to rapidly monitor the phage content of a given sample. The assay is based on the bistable expression of the *Salmonella enterica opvAB* operon. Under regular growth conditions, *opvAB* is only expressed by a small fraction of the bacterial subpopulation. In the OpvAB^ON^ subpopulation, synthesis of the OpvA and OpvB products shortens the O-antigen in the lipopolysaccharide and confers resistance to phages that use LPS as a receptor. As a consequence, the OpvAB^ON^ subpopulation is selected in the presence of such phages. Using an *opvAB::gfp* fusion, we could monitor LPS-binding phages in various media, including raw water samples. To enlarge our phage-biosensor panoply, we also developed several coliphage biosensors that proved efficient to detect LPS- as well as protein-binding coliphages. Moreover, the combination of these tools allows to identify what is the bacterial receptor triggering phage infection. The *opvAB*::*gfp* biosensor thus comes in different flavours to efficiently detect a wide range of bacteriophages and identify the type of receptor they recognize.

**Importance:** Detection and accurate counting of bacteriophages, the viruses that specifically infect bacteria, from environmental samples still constitutes a challenge for those interested in isolating and characterizing bacteriophages for ecological or biotechnological purposes. This work provides a simple and accurate method based on the bi-stable expression of genes that confer resistance to certain classes of bacteriophages in different bacterial models. It paves the way for future development of highly efficient phage biosensors that can discriminate among several receptor-binding phages and that could be declined in many more versions. In a context where phage ecology, research, and therapy are flourishing again, it becomes essential to possess simple and efficient tools for phage detection.

## Introduction

Bacteriophages, the viruses that infect bacteria, are ubiquitous on Earth, very abundant and highly diverse (1–4). They participate in the daily turnover of bacterial communities and are hypothesized to be major drivers of carbon recycling (5). Their abundance and diversity have long been acknowledged in the oceans and more recently associated to numerous microbiomes as a major part of the viromes (2, 6–8). Bacteriophages are considered a vast reservoir of genes and a major vector for horizontal gene transfer that allow the emergence of new biological functions (9, 10). Furthermore, phages have been at the origin of many discoveries and concepts in molecular biology (11–13).

Among the applications resulting from a century of phage research, phage therapy stands out by receiving more and more attention nowadays. Although experimented since the 1920s by F. d’Hérelle and subsequently abandoned in Western countries following the development of antibiotherapies (14), phage therapy has a long continuous history and usage in Eastern Europe (15). Undoubtfully, the major threat to public health represented by antibiotic resistance constitutes a breeding ground for modern phage therapy (16–18). Subsequently, many academic groups and biotech companies are hunting for more phage resources in diverse environments, thus highlighting the need for novel detection and quantification methods. Another popular application is the use of bacteriophages to specifically detect bacteria in environmental samples (19). We recently developed several versions of such biosensors able to detect enterobacteria using different readouts such as fluorescence or luminescence (20, 21).

A key aspect of bacteriophage infection, which is relevant for characterization as well as detection, is the receptor binding step. Phages recognize a variety of receptors located at the surface of bacteria including lipopolysaccharide (LPS), outer membrane proteins, pili and flagella (22–24). Extracellular appendages are used by bacteriophages to specifically recognize their target cells and irreversible binding to the appropriate surface receptor triggers genome ejection (25–28). Thus, bacteriophage binding to receptors constitutes the primary and mandatory step to a successful infection. Highlighting this importance, many studies aim at characterizing these interactions at the molecular level to understand and predict the host-range and specificity of bacteriophages.

In enterobacteria, the LPS plays an important role in host recognition by phages that belong to 2 major categories: (i) phages that use LPS as a receptor and genome ejection trigger (*eg. Salmonella* P22), and (ii) phages that bind loosely to LPS, and then need an outer membrane protein for genome injection (*eg. E. coli* T5). For both categories, different types of modification of the O-antigen can affect recognition such as the length or the glycan composition of the O-antigen (29–31). The *Salmonella enterica opvAB* operon encodes two inner membrane proteins that shorten the O-antigen chain length. Transcription of *opvAB* undergoes phase variation under the control of Dam methylation and OxyR (32). Phase-variable synthesis of OpvA and OpvB proteins results in bacterial subpopulation mixtures of standard (OpvAB^OFF^) and shorter (OpvAB^ON^) O-antigen chains in the LPS exposed at the cell surface. The dramatic change in LPS structure caused by *opvAB* expression renders *S. enterica* resistant to several bacteriophages that recognize and need the O-antigen for successful infection such as 9NA, Det7, and P22 (30, 33, 34). As a result, in the presence of these bacteriophages, the OpvAB^ON^ subpopulation is enriched, which can be monitored using an *opvAB*::*gfp* fusion (30). This interesting property was used to design a highly sensitive bacteriophage biosensor tool, allowing to detect tiny amounts of phages according to the bacterial receptor they use. We show results for two different biosensor versions able to detect phages using either the LPS or the FhuA protein as a receptor. Variants of this biosensor type could be engineered depending on the type of phage one would like to detect and quantify.

## Methods

### Bacterial strains

A full list of bacterial strains used in this study is in Table 1. Strains of *Salmonella enterica* belong to serovar Typhimurium, and derive from the mouse-virulent strain ATCC 14028. For simplicity, *Salmonella enterica* serovar Typhimurium is routinely abbreviated as *S. enterica*. Strain SV6727, harboring an *opvAB*::*gfp* transcriptional fusion downstream of the *opvB* stop codon (OpvAB^+^), was described in (30). *E. coli* K12 MG1655 was provided by the E. coli Genetic Stock Center. DR3, an MG1655 derivative with the lactose operon deleted (40), was used as an intermediate strain for the construction of DR29 and DR30. For construction of DR29, the *opvAB*::*gfp* fusion was amplified from SV6727 using the oligonucleotides MG-opvA and MG-opvB. For construction of DR30, the *opvAB::gfp* mutGATC construction was amplified from SV6729 using MG-opvA and MG-opvB. PCR products were recombined into the chromosome of DR3 at the *lac* locus (35).

**Table 1:**
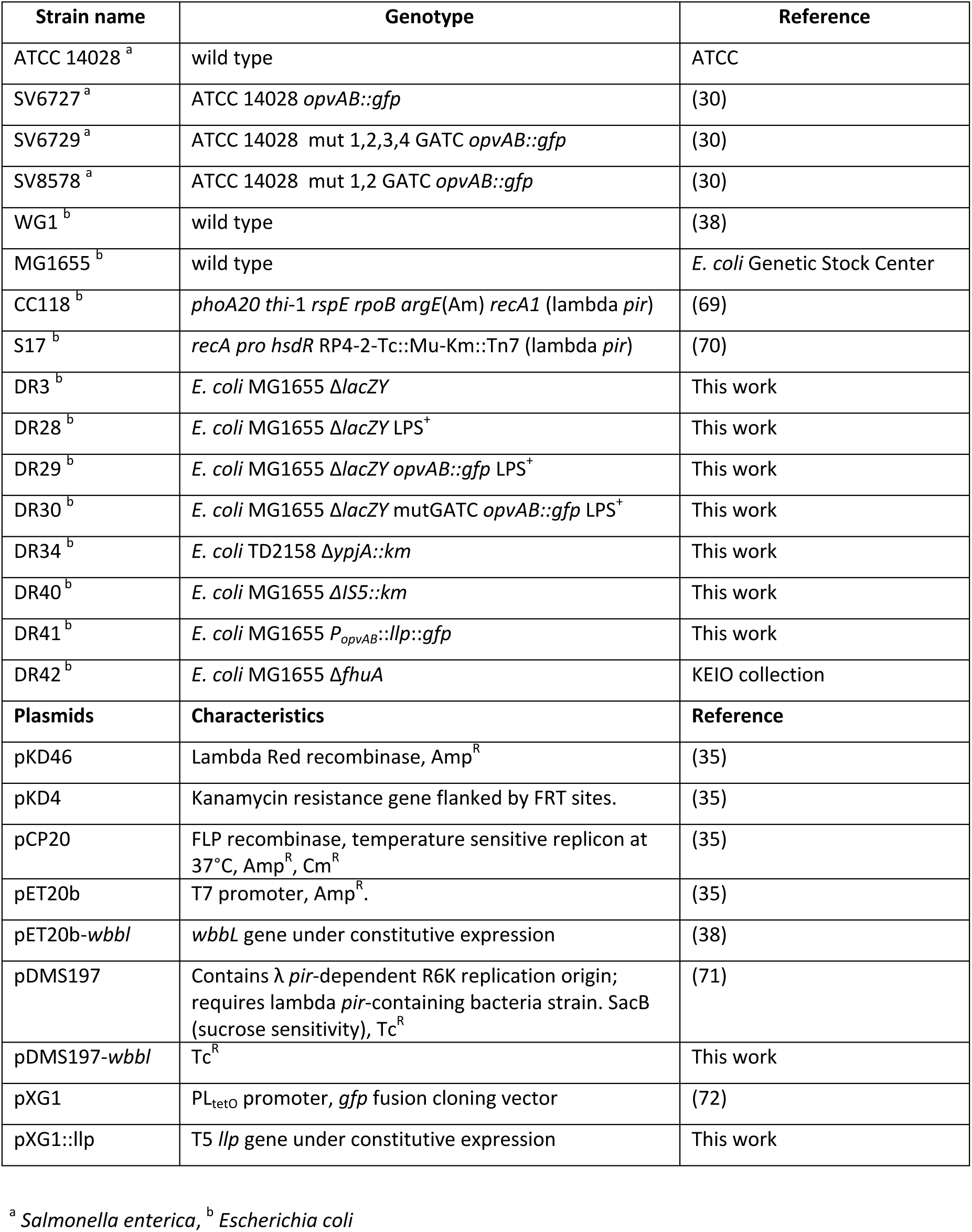
Bacterial strains and plasmids used in this study.

### Restoration of O-antigen synthesis in *E. coli* MG1655

*E. coli* MG1655 is unable to synthesize O-antigen due to a disruption of the *wbbL* gene caused by an IS5 element (36). The WbbL protein is a rhamnose transferase necessary for O-antigen synthesis (37). For complementation assays, a pETb vector containing a copy of the wild type *wbbl* gene [kindly provided by D.F. Browning (38)], was transformed into strain DR3 as a positive control of O-antigen restoration upon gene complementation. In order to replace the mutant *wbbl* gene on the MG1655 chromosome, the wild type *wbbl* gene was amplified from *E. coli* WG1 using oligonucleotides SacI-wbbl1 and XbaI-wbbl2. The resulting PCR fragment was cloned onto suicide plasmid pDMS197 (39) after restriction enzyme digestion to obtain plasmid pDMS::*wbbl*. This plasmid was propagated in *E. coli* CC118 λ *pir*. Plasmids derived from pDMS197 were transformed into *E. coli* S17 λ *pir*. The resulting strain was then used as a donor for mating with DR40 harboring a Km^R^ cassette, which was integrated into the chromosome using IS5-PS1 and IS5-PS2 oligos and pKD4 as a DNA template (35). Tc^R^ transconjugants were selected on E plates supplemented with tetracycline. Several Tc^R^ transconjugants were grown in nutrient broth (NB) and plated on NB supplemented with 5% sucrose. Individual tetracycline-sensitive segregants were then screened for kanamycin sensitivity, and the affected chromosome region was sequenced after PCR amplification with external oligonucleotides. The resulting DR28 strain is an *E. coli* MG1655 derivative with the *lac* operon deleted and the O-antigen restored.

### Construction of an *E. coli* strain expressing T5 lipoprotein

The T5 lipoprotein *llp* gene was amplified by PCR and cloned into pXG1 to obtain pXG1::Llp. For the construction of strain DR41, the T5 *llp* and *gfp* genes were amplified and linked together by a fusion PCR using opvAB-llp-F/llp-R and gfp-F/gfp-R oligos. The resulting PCR product (*llp::gfp*) was integrated at the *lac* locus of DR3. A synthetic ribosome binding site named BI-RBS used in a previous study (40) was inserted upstream of the *llp* gene to optimize Llp production.

### Bacteriophages

Bacteriophages 9NA (34, 41) and Det7 (33) were kindly provided by Sherwood Casjens, University of Utah, Salt Lake City. Bacteriophage P22_H5 is a virulent derivative of bacteriophage P22 that carries a mutation in the *c2* gene (42), and was kindly provided by John R. Roth, University of California, Davis. T5 bacteriophage was provided by Pascale Boulanger, I2BC, Orsay, France. Bacteriophages Se_F1, Se_F2, Se_F3 and Se_F6 infecting *S. enterica* were isolated and purified from waste water samples in Seville. Genome sequences are under submission. Other waste water bacteriophages were isolated upon infection of either DR28 (*E. coli* MG1655 Δ*lacZY* LPS^+^) or MG1655. A list of all the phages isolated in this study is included in Table 3.

**Table 2:**
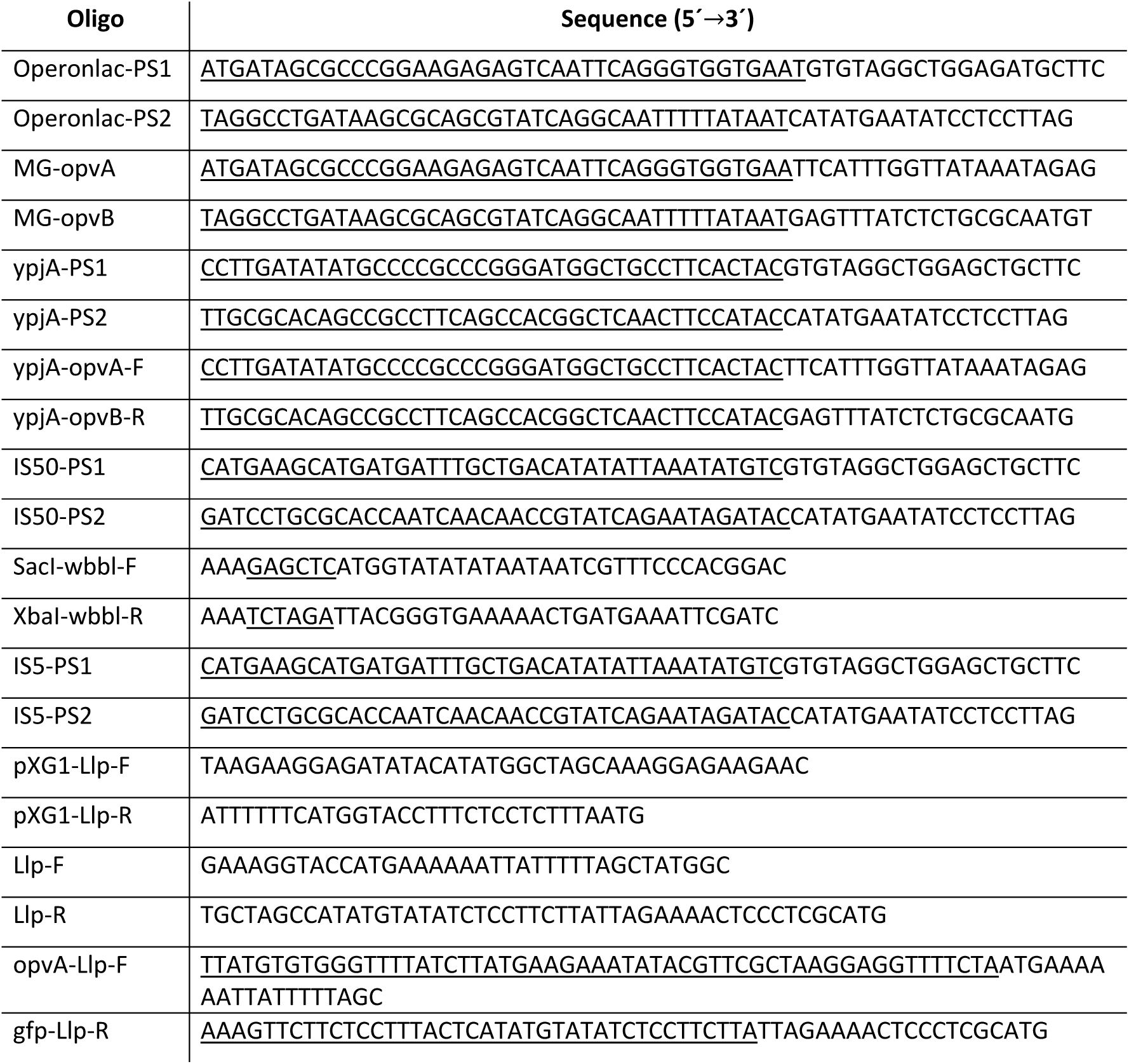
Oligonucleotides used in this study.

**Table 3.** Bacteriophages used in this study.

| Bacteriophage | Host | Origin | Reference |
| --- | --- | --- | --- |
| P22 | *S. enterica* | Sherwood Casjens |  |
| Det7 | *S. enterica* | Sherwood Casjens |  |
| 9NA | *S. enterica* | Sherwood Casjens |  |
| T5 | *E. coli* MG1655 | Pascale Boulanger |  |
| Se_F1 | *S. enterica* | Sevilla, Spain (Waste water) | This work |
| Se_F2 | *S. enterica* | Sevilla, Spain (Waste water) | This work |
| Se_F3 | *S. enterica* | Sevilla, Spain (Waste water) | This work |
| Se_F6 | *S. enterica* | Sevilla, Spain (Waste water) | This work |
| Se_ML1 | *S. enterica* | Sevilla, Spain (Maria Luisa Park) | This work |
| Se_AO1 | *S. enterica* | Sevilla, Spain (Sheep farm) | This work |
| Ec_Unk_EM | DR28 | Sevilla, Spain (Waste water) | This work |
| Ec_Unk_EM1 | DR28 | Sevilla, Spain (Waste water) | This work |
| Ec_Unk_EM2 | DR28 | Sevilla, Spain (Waste water) | This work |
| Ec_Unk_AO1 | DR28 | Sevilla, Spain (Sheep farm) | This work |
| Ec_Unk_ML | DR28 | Sevilla, Spain (Maria Luisa Park) | This work |
| Ec_Unk_ML1 | DR28 | Sevilla, Spain (Maria Luisa Park) | This work |
| Ec_Unk_MLB | DR28 | Sevilla, Spain (Maria Luisa Park) | This work |
| Ec_Unk_PO1 | MG1655 | Cáceres, Spain (chicken farm) | This work |
| Ec_Unk_PO2 | MG1655 | Cáceres, Spain (chicken farm) | This work |
| Ec_Unk_PO3 | MG1655 | Cáceres, Spain (chicken farm) | This work |

### Culture media

Bertani’s lysogeny broth (LB) was used as standard liquid medium. Nutrient broth (NB) was used to recover cultures after transduction or transformation. Solid media contained agar at 1.5% final concentration. Green plates (43) contained methyl blue (Sigma-Aldrich, St. Louis, MO) instead of aniline blue. Antibiotics were used at the concentrations described previously (44). E50X salts [H_3_C_6_H_5_O_7_ · H_2_O (300 g/l), MgSO_4_ (14 g/l), K_2_HPO_4_ · 3H_2_O (1965 g/l), NaNH_4_HPO_4_ · H_2_O (525 g/l)] were used to prepare E minimal plates [E50× (20ml/l), glucose (0.2%), agar 20 g/l].

### Electrophoretic visualization of LPS profiles

To investigate LPS profiles, bacterial cultures were grown overnight in LB. Bacterial cells were harvested and washed with 0.9% NaCl. Around 3×10^8^ cells were pelleted by centrifugation. Treatments applied to the bacterial pellet such as electrophoresis of crude bacterial extracts and silver staining were performed as described elsewhere (45).

### Bacteriophage challenge

Bacterial cultures were grown at 37°C in LB (5 ml) containing phages (100 μl of phage lysate [10^8^-10^10^pfu]). Cultures were diluted 1:100 in LB + phage and incubated until exponential phase (OD_600_ ∼ 0.3) before flow cytometry analysis. To monitor bacterial growth, overnight bacterial cultures were diluted 1:100 in 200 μl of LB. Five μl of a bacteriophage lysate were added (10^8^-10^10^ pfu). OD_600_ and fluorescence intensity were subsequently measured at 30 min intervals using a Synergy™ HTX Multi-Mode Microplate Reader from Biotek.

### Flow cytometry

Bacterial cultures were grown at 37°C in LB or LB containing phage until exponential (OD_600_ ∼0.3) or stationary phase (OD_600_ ∼4). Cells were then diluted in PBS to obtain a final concentration of ∼10^7^ cfu/ml. Data acquisition and analysis were performed using a Cytomics FC500-MPL cytometer (Beckman Coulter, Brea, CA). Data were collected for 100,000 events per sample, and analyzed with CXP and FlowJo 8.7 software.

### Phage isolation from water samples

Samples of water were filtered using a Millex-GS filter, 0.22 μm filter pore. Each sample was divided into two samples; chloroform was added to one of the parts. The filtered sample of water (1 ml) was added to an overnight culture of host bacterial strain (0.5 ml). The mixture was supplemented with 3.5 ml of LB broth and was incubated overnight at 37℃ without shaking. The culture was then centrifuged at 5000 rpm during 5 min. The supernatant was filtered using a filter, 0.22 μm pore size, in order to discard bacterial cells and debris. 50 μl of an overnight culture of the bacterial target was spread on LB plates. Lysates were dropped onto the agar surface, and left to dry. The plates were inspected for lysis zones after overnight incubation at 37°C. The spot assay was used to assess the bactericidal ability of different phage lysates. Isolation of phage was performed by the double agar layer method using *Salmonella* and *E. coli* as a host system. Isolated plaques were suspended in LB and were streaked in plates which contained a bacterial layer in order to isolate pure phage. Phages were characterized on the basis of plaque morphology and host range. Moreover, certain phages were characterized by electron microscopy and their genomes were sequenced.

### Electron microscopy

Electron microscopy was performed at the Mediterranean Institute of Microbiology (IMM) facility. Phage suspensions (5 µl) were dropped onto a copper grid covered with Formvar and carbon and left 3 minutes at 25°C, after what the excess of liquid was removed. Phage particles were stained according to (46) with a 2% uranyl acetate solution. Dried grids were then observed using a transmission microscope FEI Tecnai 200 kV coupled to an Eagle CCS 2kx2k camera. Images were then analyzed using the ImageJ software.

### Detection of phage in a crude water sample

A tube with 3.5 ml of LB, 0.5 ml of an overnight culture of SV6727 strain (*opvAB*::*gfp*) and 1 ml of a 0.22 µm filtered water sample was incubated for 7 h without shaking. The culture was diluted at a ratio of 1/200 in LB medium and incubated at 37°C with shaking (200 rpm) for 3 hours before dilution in PBS buffer. All water samples were collected in the Sevilla area. Data acquisition and analysis were performed using a Cytomics FC500-MPL cytometer (Beckman Coulter, Brea, CA). Data were collected for 100,000 events per sample, and analyzed with CXP and FlowJo 8.7 software. The presence of phages in water samples was checked simultaneously by plating 1 ml of water on a plate with *S. enterica* poured into soft agar.

## Results

### Proof of concept using characterized bacteriophages

Based on results published by Cota and collaborators (30), we foresaw that the *opvAB* phase variation operon could be used as a bacteriophage biosensor. Indeed, the OpvAB^OFF^ subpopulation was shown to be killed by bacteriophages that belonged to different families and used LPS for infection, leading to enrichment of the OpvAB^ON^ subpopulation. This population is insensitive to such infections due to the shortening of its LPS by the *opvAB* gene products (30). Thus, if an *opvAB::gfp* fusion was used, enrichment of OpvAB^ON^ cells could be detected by increased fluorescence intensity (30). To start with, we wanted to compare the efficiency of two different methods of fluorescence detection, flow cytometry and plate reader monitoring, to detect 3 different bacteriophages known to use LPS for infection. As shown in Figure 1A, flow cytometry turned out to be not only very sensitive but also descriptive of the bacterial population heterogeneity. In contrast, fluorescence detection using a plate reader only provided a rough assessment of the population structure (Fig. 1B). On the other hand, fluorescence detection using a plate reader does not necessitate a high-cost machine such as a flow cytometer and has the advantage to give a measure according to the OD_600_ of the culture. Our standard protocol for phage detection by flow cytometry includes 8 h of incubation to ensure that phages kill the vast majority of OpvAB^OFF^cells, thus leading to the enrichment of the OpvAB^ON^ population. However, thanks to the sensitivity of this technique and because fluorescence increases as a function of time, measurements can be done earlier than 8 h. As depicted in Figure 1C, incubation for 1 or 2 h after dilution into fresh medium in the presence of P22_H5 led to 2.6% and 11.3% of positive cells respectively. Since flow cytometry is highly reproducible, these levels are sufficient to discriminate an OpvAB^ON^ subpopulation from the background level of cells that passed the fluorescence threshold in the absence of phage (0.2%) (Fig. 1C). These experiments thus validated the proof-of-concept of a fluorescent biosensor for the detection of LPS-using bacteriophages based on a modified *opvAB* operon.

**Figure 1.**
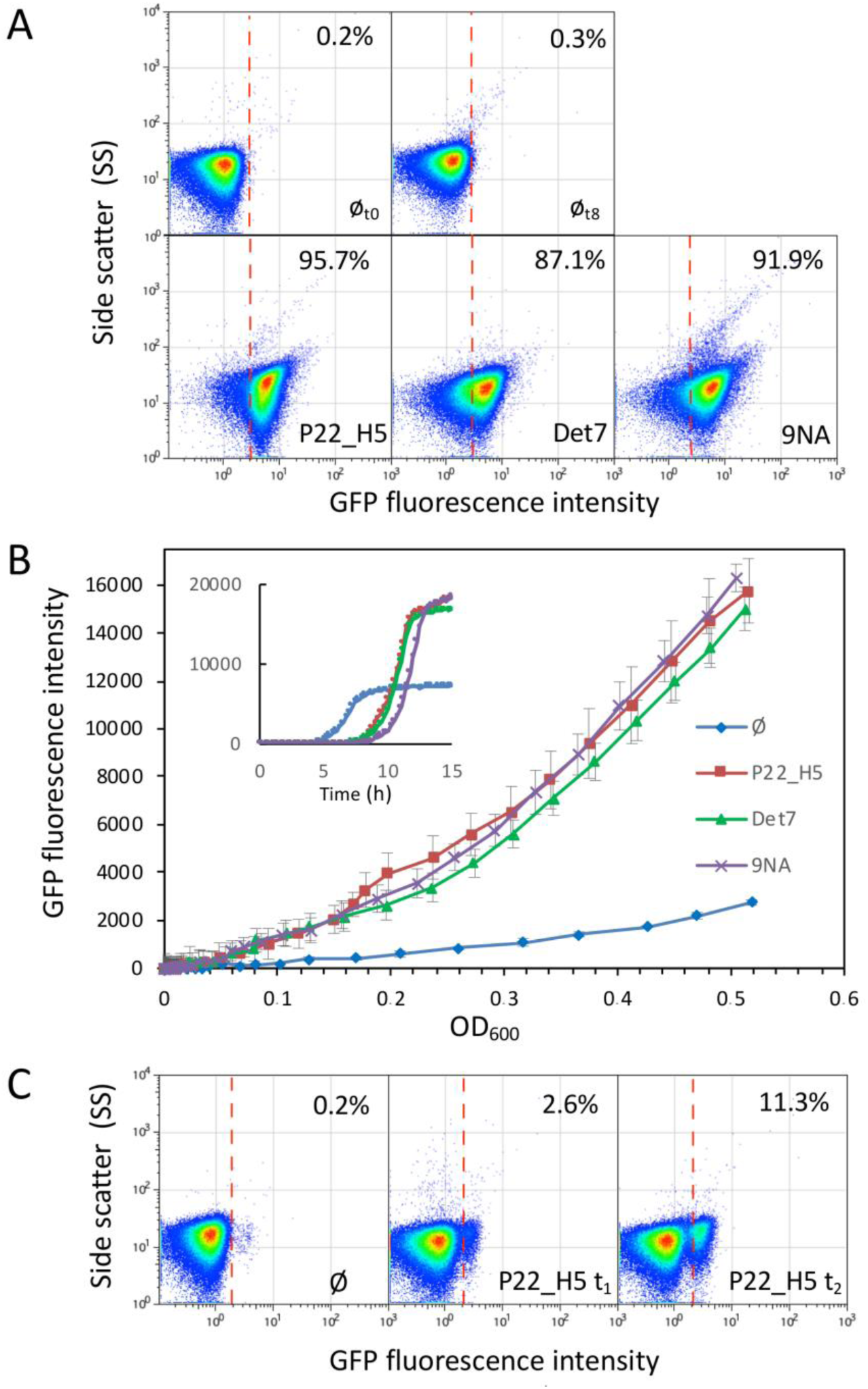
Detection of increased fluorescence intensity upon selection of the OpvAB^ON^ subpopulation in the presence of bacteriophages. **A.** GFP fluorescence distribution in an *S. enterica* strain carrying an *opvAB::gfp* fusion (SV6727) before (t = 0 h) and after growth in LB without phage, or in the presence of either P22 H5, 9NA, or Det7 (t = 8 h). Data are represented by a dot plot (side scatter versus fluorescence intensity [ON subpopulation size]). All data were collected for 100,000 events per sample. **B.** Growth curves of strain SV6727 in contact with P22 H5, Det7 or 9NA phage. Data are represented by growth curves [Fluorescence intensity] versus growth [OD_600_] or [time] (insert). **C.** GFP fluorescence distribution of strain SV6727 before (t = 0 h) and after growth in LB containing Det7 (t = 1 h or 2 h).

### Detection of uncharacterized bacteriophages

To go further with the phage detection tool, we decided to isolate and purify various *S. enterica*-infecting bacteriophages and test whether they could be detected without previous characterization of the phage receptor. Indeed, a phage that would need a functional and extended LPS should kill the OpvAB^OFF^ subpopulation as the known phages did. In contrast, a phage that would not need (or that can bypass) LPS to infect should not discriminate the two sub-populations. Various phages were enriched, isolated and partially characterized from different environments in the Seville area as described in the Methods section. Without going deeper into the characterization of those phages, the plaque morphologies (not shown), and even more the images obtained after negative staining by TEM indicated that the 6 isolated phages were different from each other (Fig. 2A). This was further confirmed by sequence determination of the 6 phage genomes (Olivenza DR et al, in preparation). The 6 of them were assayed by flow cytometry for detection using the *opvAB::gfp* fusion. Interestingly, all 6 were perfectly detected 8 hours post-infection with more than 60% of the cells in the ON state (Fig. 2B). This proportion reached up to 90 % for three individual phages (Se_F1, Se_F6 and Se_AO). In contrast, Se_F3 and Se_ML killed the OpvAB^OFF^ subpopulation only partially. A conclusion from these experiments was that the *opvAB::gfp* fusion provides an efficient tool to detect unknown LPS-dependent bacteriophages of *S. enterica*. The variations observed between the different flow cytometry profiles could be due to a difference in the usage of the LPS as a receptor for those phages or to a partial resistance of the OFF supopultion that could prevent OpvAB^ON^ subpopulation enrichment.

**Figure 2.**
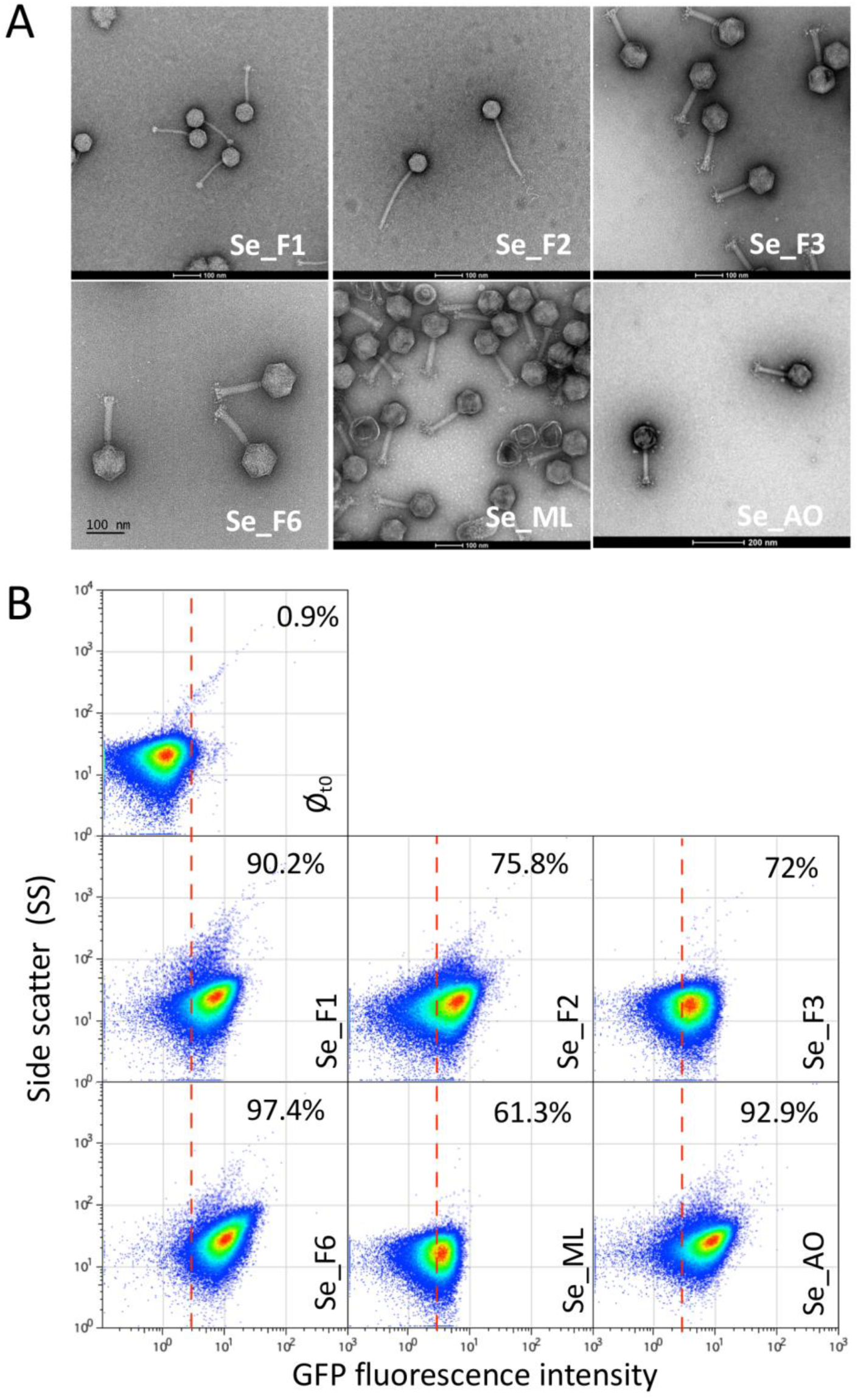
Detection of new bacteriophages. **A.** GFP fluorescence distribution in strain SV6727 (*opvAB::gfp*) before (t = 0) and after growth in LB containing new purified bacteriophages (Se_F1, Se_F2, Se_F3, Se_F6, Se_ML and Se_AO) after 8 h. Data are represented by a dot plot (side scatter versus fluorescence intensity [ON subpopulation size]). The percentage of ON cells in each sample is indicated. All data were collected for 100,000 events per sample. **B.** Negative staining of bacteriophages visualized by electron microscopy.

An especially appealing objective of the present tool is to use *opvAB::gfp* fusion to detect bacteriophages in crude samples. Different crude water samples collected in the Seville area were assayed for the presence of phages in the presence of the biosensor strain for 10 h before analyzing the resulting populations by flow cytometry. Among the four samples tested, three (EM1, EM2 and EM3) displayed a significant increase in GFP activity, indicating that indeed these samples contained bacteriophages depending on LPS for infection (Fig. 3). This conclusion was confirmed by plaque assay (not shown). One sample (AO) did not show increase of the OpvAB^ON^ subpopulation, which suggested that the sample did not contain any LPS-depending phage active against *S. enterica*. Alternative explanations were that the sample might contain a phage inhibitor, which is unlikely since we obtained plaques on a *S. enterica* lawn, or that phages present in the sample infected both OpvAB^ON^ and OpvAB^OFF^ *S. enterica*. Taken together these results prove that our *opvAB*-based fluorescent biosensor can efficiently detect the presence of LPS-depending bacteriophages in environmental samples without any pre-processing other than incubation with the biosensor.

**Figure 3.**
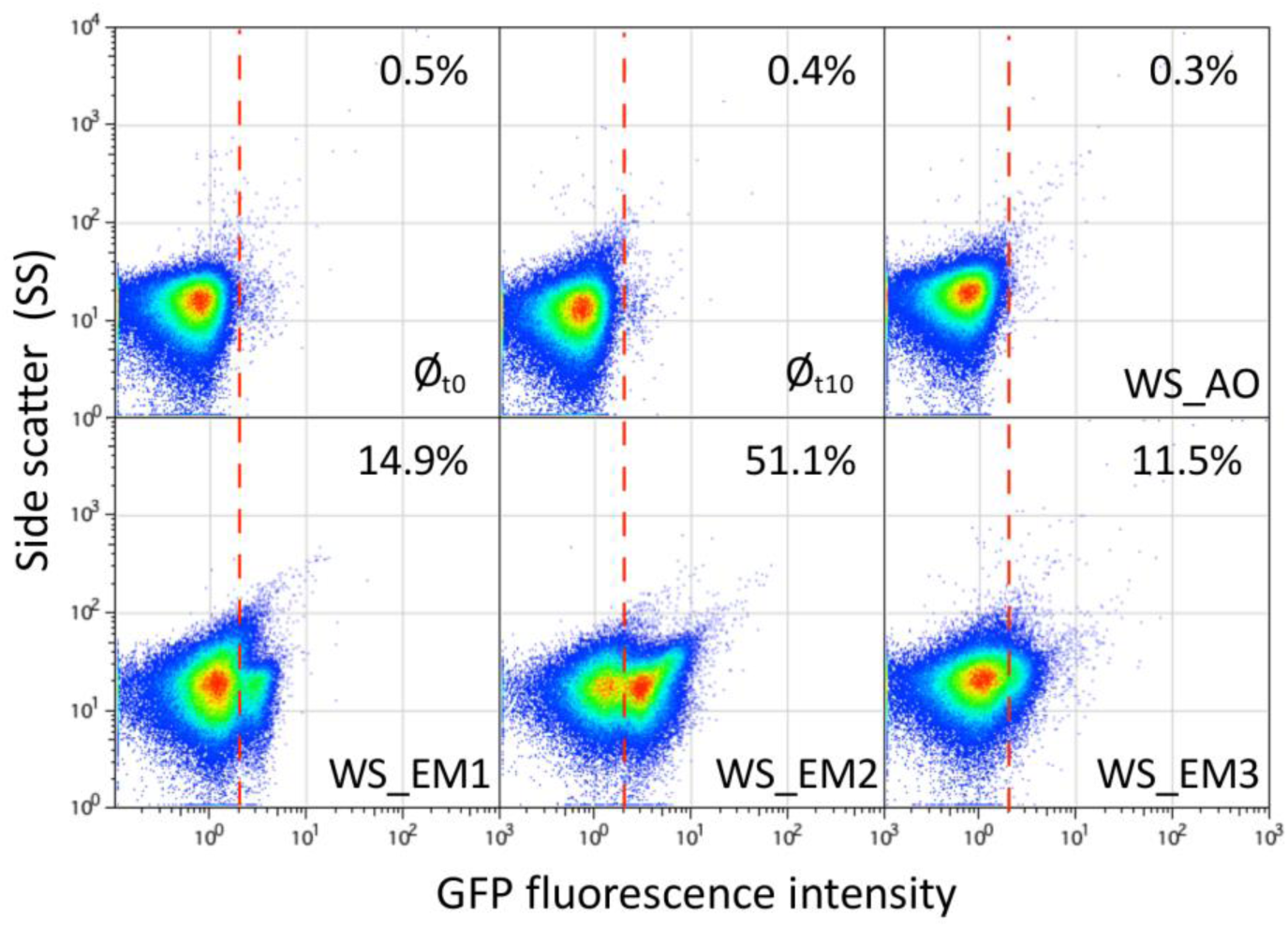
Detection of bacteriophages from crude water samples. GFP fluorescence distribution in strain SV6727 (*opvAB::gfp*) before and after growth in LB for 10 hours in the presence of crude water previously filtered. Data are represented by a dot plot (side scatter versus fluorescence intensity [ON subpopulation size]). The ON subpopulation percentage is shown for each sample. Data were collected for 100,000 events per sample. The presence of bacteriophages was previously verified by a plaque assay (20-30 pfu/ml) on a *S. enterica* lawn.

### Sensitivity and optimization of the detection tool

To challenge the limits of the phage detection tool, 9NA was used (Fig. 1). We first determined the detection limit in a fluorescence plate reader by using serial dilutions of 9NA. As shown in Fig. 4A, only the most diluted sample, equaling to 10 PFU/ml, could not be distinguished from the control experiment with no phage. Therefore, using a fluorescence plate reader, as little as 100 PFU/ml could be detected. Interestingly, the curves obtained in the presence of effective amounts of phage show an interesting profile. In the first five hours, the global tendency was an increase of the whole population (OpvAB^ON^ and OpvAB^OFF^), followed by a rapid decrease (Fig. 4A, inset). Five hours post-infection, the bacterial growth resumed more or less rapidly depending on the amount of phage initially present in the culture. As expected, the fluorescence intensity correlated with growth only when samples were initially infected with 9NA (≥ 100 PFU/ml) (Fig. 4A), indicating that only the ON subpopulation grew under these conditions. This was confirmed by the fact that the highest fluorescence intensity was reached in the presence of the highest amount of phage, which killed the OpvAB^OFF^ subpopulation more efficiently. As expected, in the absence of phage (or in the presence of a low number of phages), bacterial growth did not correlate with fluorescence increase, indicating that the OpvAB^OFF^ population mainly accounts for bacterial growth. Based on this experiment, we tentatively concluded that strong, dose-dependent fluorescence intensity was reached 10 hours post infection (Fig. 4A, inset). In order to test those parameters using flow cytometry, the same serial dilutions of phages were applied for 10 h and enrichment of the OpvAB^ON^ subpopulation was monitored. In contrast with the plate reader experiment, all the phage concentrations tested above 1 PFU/ml were able to enrich the ON subpopulation (Fig. 4B). Moreover, a linear correlation was observed between the phage concentration and the OpvAB^ON^ subpopulation size up to 10^2^ PFU/ml, which could be used as a calibration curve to determine phage concentration in a given sample. Hence, the phage biosensor appears to be highly sensitive and can detect as little as 10 PFU/ml.

**Figure 4.**
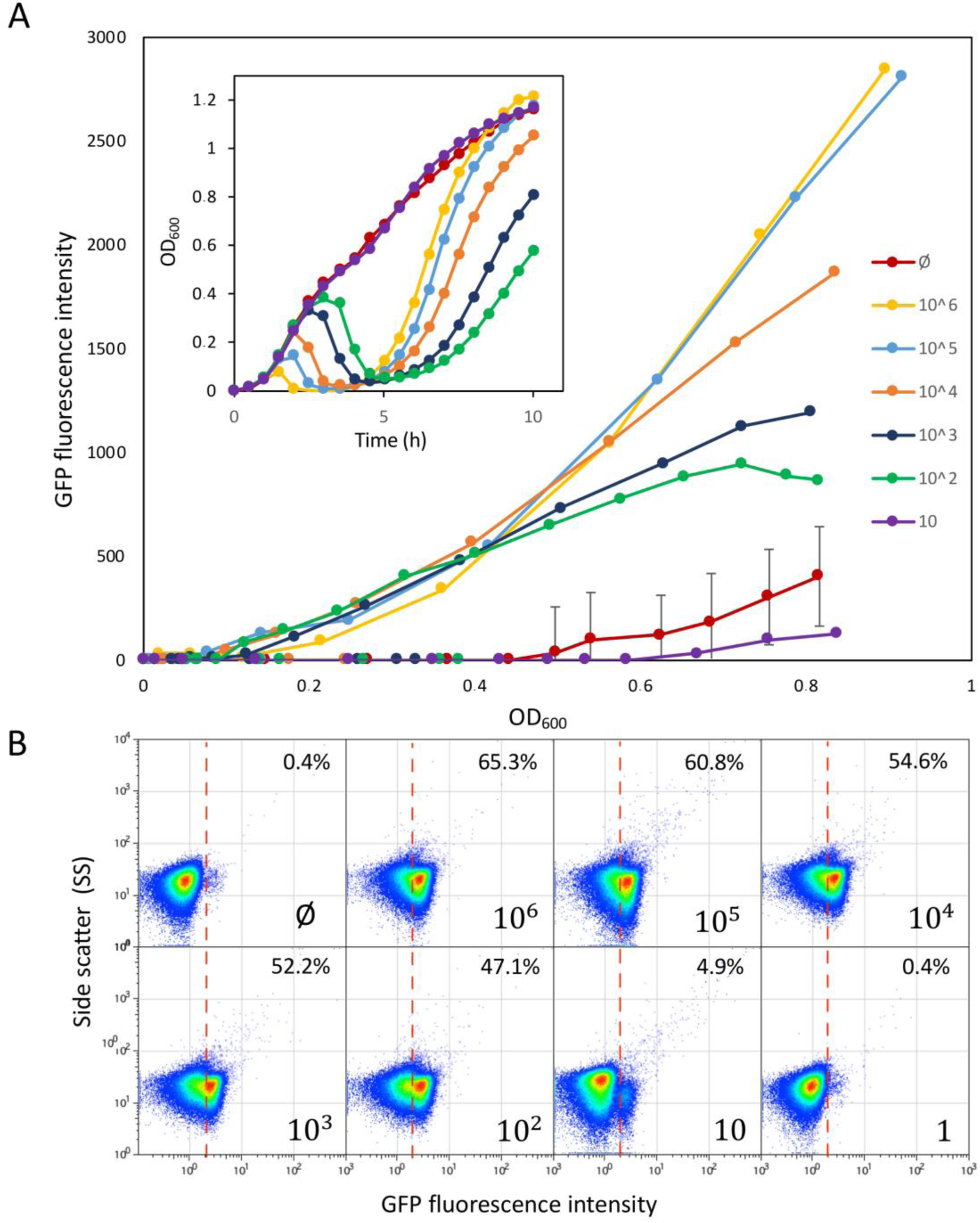
OpvAB phage-biosensor sensitivity. **A.** Growth curves of SV6727 (*opvAB::gfp*) in contact with serial dilutions of phage 9NA, from 10^6^ up to 10^1^ PFU/ml. Data are represented by growth curves showing OD_600_ versus time [h] (inset), and (fluorescence intensity [ON subpopulation size] versus growth [OD_600_]. **B.** GFP fluorescence distribution in SV6727 before (t = 0) and after growth (t = 10 h) in LB containing serial dilutions of 9NA phage starting from 10^6^ up to 10^0^ PFU/ml. Data are represented by a dot plot (side scatter versus fluorescence intensity [ON subpopulation size]). All data were collected for 100,000 events per sample.

Our next objective was to improve the detection limits of our biosensor. As we noticed that killing of the OpvAB^OFF^ subpopulation (and thus selection of the OpvAB^ON^ subpopulation) was more efficient in the presence of a high number of 9NA PFU (Fig. 4A), we aimed at improving the method by adding less cells to increase the ratio phage / biosensor. Indeed, the more diluted were the cells, the smaller number of phages that could be detected (Fig. 5A). It is remarkable that for the smallest dilution of the overnight culture the concentration of phages detected using a microplate reader was around 4.10^5^ PFU/ml, whereas as little as 4 PFU/ml were detected using a higher dilution of the biosensor (equivalent to 8.8×10^4^ cells/ml, corresponding to about 17,600 ON cells), thus increasing 5-log the detection limit when the number of cells decreased 3-log only.

**Figure 5.**
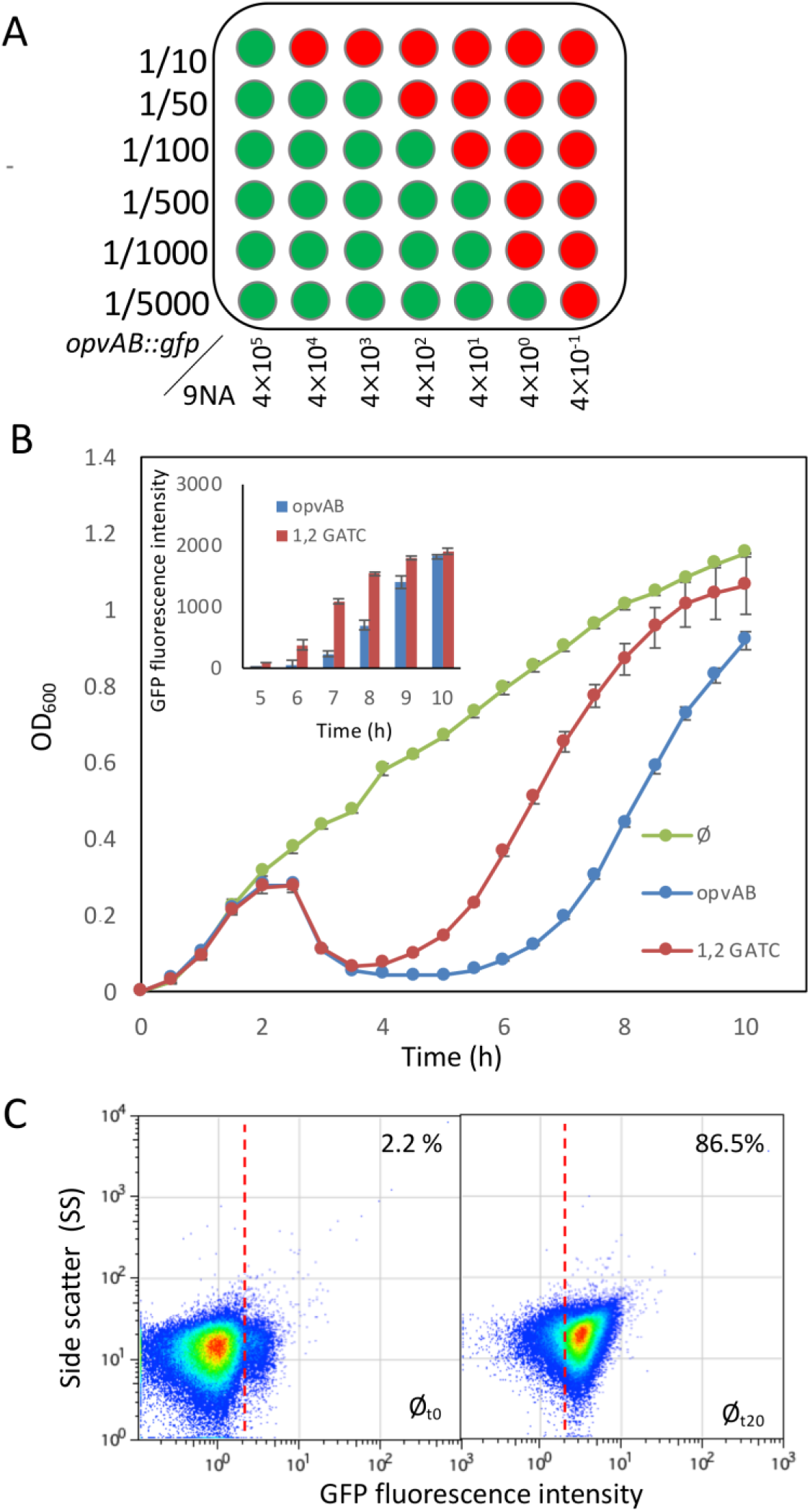
Improving the *opvAB* biosensor sensitivity. **A.** Bacterial dilutions (10^-1^ up to 2.10^-4^) of strain SV6727 grown in the presence of serial dilutions of bacteriophage 9NA. Bacterial growth was monitored in a microplate reader. Green wells are fluorescence-positive, and red wells are fluorescence-negative. **B.** Growth curves obtained for a strain expressing the *gfp* fusion under the control of a wild type promoter (SV6727) in the absence of phage (green) and in the presence of 9NA (blue), and for a derivative strain carrying point mutations in two GATC sites at the *opvAB* promoter region (SV8578) in the presence of phage 9NA (red). Fluorescence intensity data were obtained at several time points of the growth curve (inclusion). **C.** GFP fluorescence distribution in SV8578 (GATC_1,2_ *opvAB::gfp)* before (t = 0 h) and after growth in LB containing 9NA phage (t = 20 h). Data are represented by a dot plot (side scatter versus fluorescence intensity [ON subpopulation size]). All data were collected for 100,000 events per sample.

Another improvement originated from earlier work showing that the OpvAB^ON^ sub-population size increased when specific GATC sites present in the promoter region were mutated (30, 32). Indeed, when the mut1,2GATC *opvAB::gfp* construct was used as a biosensor for phage 9NA, the fluorescence increased faster than with the construct bearing the wild-type promoter and reached a plateau in about 10 h (Fig. 5B, inset). Interestingly, the largest difference was observed around 7 h post-infection.

We then combined both improvements (low cell density and mutations in GATC sites) to reach an even lower detection limit. The experiment described in Fig. 5C was performed in two steps. First, we used as little as eight 9NA phages diluted into 500 ml of LB containing a dilution (1/10) from an overnight culture of the biosensor strain and let the phage enrichment proceed for 15 h. Then the mixture was diluted again into fresh medium to let the OpvAB^ON^ subpopulation enrich for 5 h before the flow cytometry assay. It is remarkable that an extremely small initial number of phages was able to enrich the ON subpopulation up to 86.5% under the conditions described (Fig. 5C), thus lowering the detection level to about 1.6 phages per 100 ml. As a reminder, before optimization the detection limit with flow cytometry was around 1000 phages per 100 ml.

### Design of a biosensor able to detect coliphages using the *opvAB*::*gfp* fusion

Up to this point, the design and improvement of the phage biodetection method was performed using *S. enterica* as a chassis. To widen our capacity to detect phages in environmental samples, we decided to adapt the biosensor to coliphages. To do so, we first had to restore a full length LPS as *E. coli* K12 MG1655 is well-known to carry an IS5-interrupted version of the *wbbL* gene encoding a rhamnosyltranferase (38). This is clearly evidenced by the length and profile of the LPS following extraction, separation by SDS-PAGE and silver staining is affected in MG1655 (Fig. 6A, lane 1). The interrupted *wbbL* gene was complemented either by adding a plasmid-based copy of *wbbL* or by integrating ectopically a single copy of the wild type gene. In both cases, the LPS structure was restored (Fig. 6A, lanes 4 and 5). Engineering of strains carrying an *opvAB::gfp* fusion on the chromosome allowed us to examine the consequences of *opvAB* expression in *E. coli*. A wild type *opvAB* control region caused a subtle alteration of the LPS profile, an observation consistent with the small size of the ON subpopulation (Fig. 6A, lane 6). Modification of the length distribution of glycan chains in the O antigen was however unambiguous when the LPS^+^ *E. coli* strain harbored a GATC-less *opvAB::gfp* fusion (Fig. 6A, lanes 6 and 7).

**Figure 6.**
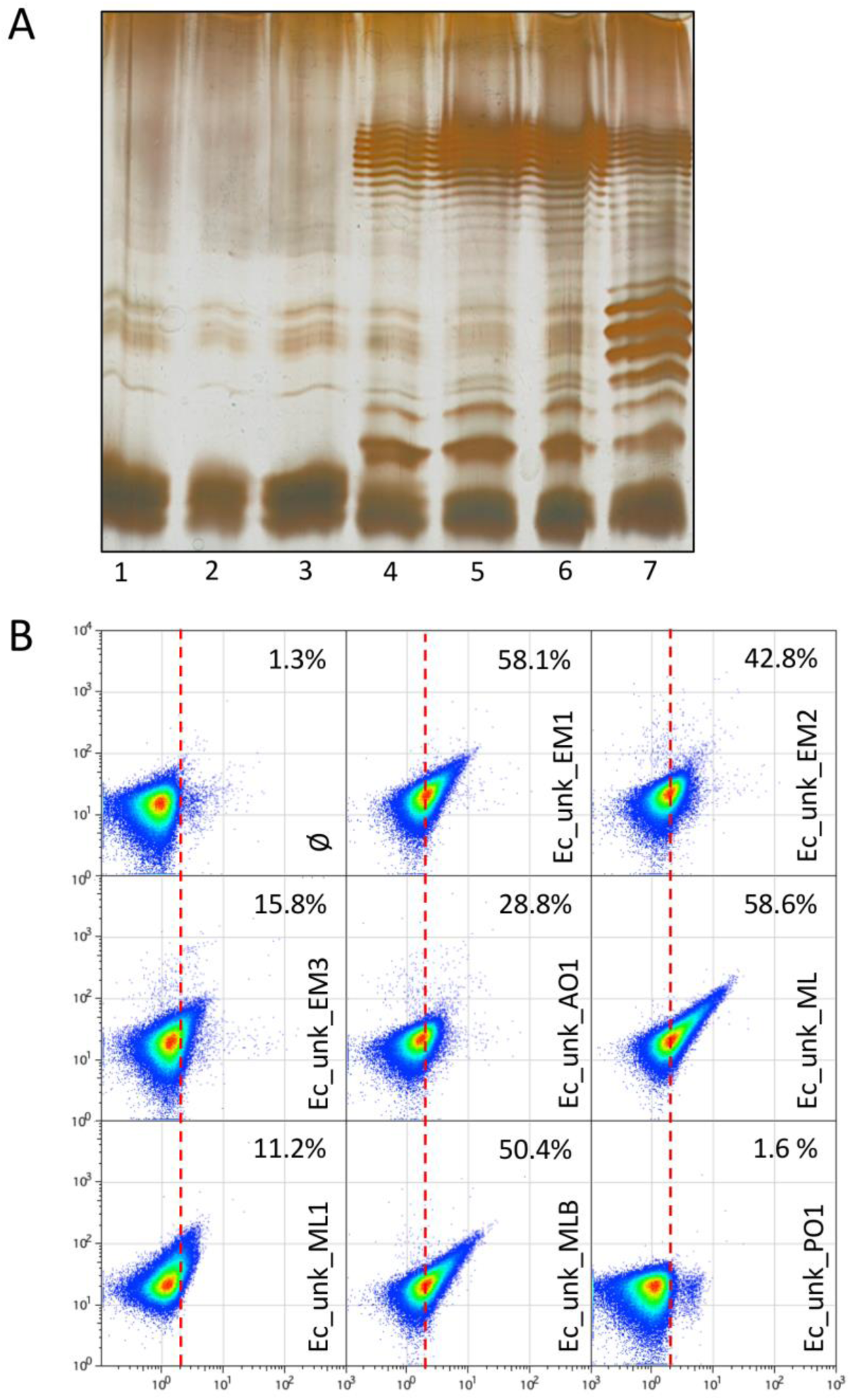
Restoration of *E. coli* MG1655 O-Antigen and detection of unknown coliphages. **A.** Electrophoretic visualization of LPS profiles from different MG1655 derivatives, as follows**: 1,** Wild type *E. coli* MG1655 strain containing an IS5-inactivated version of the *wbbl* gene; **2,** *E. coli* MG1655 with a deletion of the lactose operon (DR3); **3**, *E. coli* DR3 carrying the empty pETb vector; **4**, *E. coli* DR3 carrying a pETb derivative containing a wild type *wbbL* gene; **5**, *E. coli* DR3 carrying the *wbbL* gene integrated into the genome, replacing the altered IS5-*wbbl* gene (LPS^+^ strain); **6**, *E. coli* MG1655 LPS^+^ carrying the *opvAB::gfp* construction (DR29); **7**, *E. coli* MG1655 LPS^+^ *opvAB::gfp* GATC-less (DR30). **B.** GFP fluorescence distribution in *E. coli* DR29 before (t = 0 h) and after growth in LB, or LB containing purified uncharacterized bacteriophages (Ec_Unk_EM1, Ec_Unk_EM2, Ec_Unk_EM3, Ec_Unk_AO1, Ec_Unk_ML, Ec_Unk_ML1, Ec_Unk_MLB and Ec_Unk_PO1 (t = 8 h). Data are represented by a dot plot (side scatter versus fluorescence intensity [ON subpopulation size]). ON subpopulation sizes (percentages) are shown for each sample. All data were collected for 100,000 events per sample.

Using the *opvAB::gfp*-carrying LPS^+^ *E. coli* strain, we isolated and purified 7 coliphages from water samples from the Seville area. As shown in Figure 6B, seven of the coliphages were able to select the OpvAB^ON^ subpopulation with an enrichment efficacy ranging from 11.2 to 58.6 %, far higher than the control without phage (1.27 %). Again, this difference could account to the use of the LPS as a primary receptor or as a full receptor for the isolated coliphages except for Ec_unk_PO1. Whatever the case, this experiment shows that the method works alike in *E. coli* and in *S. enterica,* and that the *opvAB*::*gfp* fusion is a versatile tool that could be used with different enterobacterial chassis.

### Expanding the *opvAB*::*gfp* biosensor to the detection of phages that use proteins as receptors: Bacteriophage T5 detection as a proof-of-concept

As we have seen that coliphage detection could show some variation (Fig. 6B), probably depending on the receptor molecule used for recognition at the surface of the bacteria, we decided to design an *E. coli* biosensor that would specifically detect a phage known to bind a protein. Phage T5 is an appropriate model: although it uses LPS for initial adsorption, binding to the ferrichrome transporter FhuA on the outer membrane constitutes the trigger for DNA injection (27, 47). We first thought to make a simple P*_opvAB_*::*fhuA*-*gfp* fusion, but that would have provided us with a biosensor working the opposite way from the *opvAB*::*gfp* biosensor. Indeed, in this case the presence of T5 would select for the OFF (non-fluorescent) subpopulation, and it would mean we should be able to detect a slight decrease in fluorescence, which can be expected to be less reliable than monitoring an increase of fluorescence intensity. We thus looked for a gene which under the control of the *opvAB* promoter would confer resistance to T5 in the ON state. Interestingly, T5 encodes a lipoprotein, encoded by the *llp* gene, which is an inhibitor of T5 infection (48). Llp is synthesized upon T5 infection and prevents superinfection of the host by other T5 virions by interacting with the FhuA receptor, resulting in its inactivation (49). Moreover, the *llp* phage gene is expressed in the early stage of T5 infection, which not only prevents superinfection but also protects progeny phages from being inactivated by the receptor present in envelope fragments of lysed host cells (50). We thus reasoned that placing the *llp* gene under the control of the *opvAB* promoter might confer resistance to T5 only in the ON state. As expected from the literature, a Δ*fhuA* strain proved resistant to T5, but not to T4, which uses OmpC as receptor (51) (Fig. 7A). In turn, ectopic expression of *llp* from a plasmid also confers resistance to the *E. coli* MG1655 WT strain as in the Δ*fhuA* mutant. We thus integrated a P*_opvAB_*::*llp*-*gfp* fusion at the *lac* locus in *E. coli* MG1655 and assayed this new biosensor with T5 for 10 hours on a plate reader, periodically monitoring the OD_600_ and the florescence intensity (Fig. 7B). As predicted, enrichment of the ON subpopulation was detected after a few (4) hours and GFP fluorescence intensity increased, indicating that this fusion performed well as a T5 biosensor. We then assayed the biosensor using flow cytometry to estimate the efficiency of ON subpopulation enrichment, which ranged from 82% to almost 90% in 10 h for the 4 different phages isolated on *E. coli* MG1655 (not complemented) (Fig. 7C). Interestingly, addition of colicin M led to a similar enrichment of the ON subpopulation, thus confirming that production of the FhuA inhibitor Llp was also able to prevent colicin M binding. We also performed an additional control, using one of the LPS-binding phages assayed in Figure 6. As predicted based on its effect on the Ec_*opvAB*::*gfp* biosensor, this specific phage (Ec_unk_EM1) was unable to select for the P*_opvAB_*::*llp*-*gfp* ON subpopulation (Fig. 7C). Conversely, Ec_unk_PO1 (selected to turn on P*_opvAB_*::*llp*-*gfp* fusion, and thus likely recognizing FhuA) was not detected using the Ec_*opvAB*::*gfp* biosensor (Fig. 6C). Together, these experiments show that the method can accommodate not only different bacterial chassis (*Salmonella* and *E. coli*) but also different types of receptors, thus providing primary indication on the type of receptor used by a given phage.

**Figure 7.**
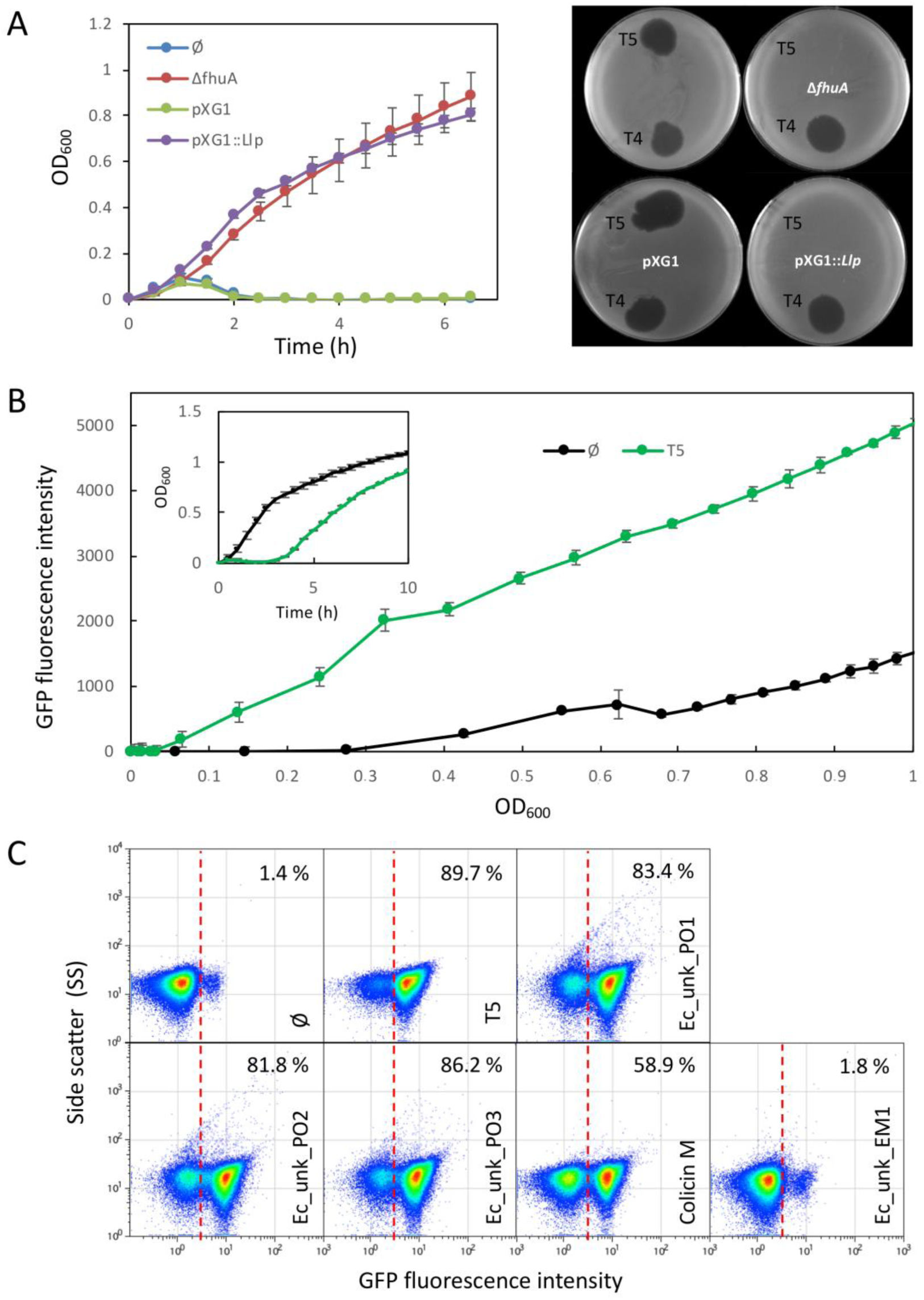
Bacteriophage T5 detection. **A. Left**. Bacterial growth curves in the presence of phage T5 (2.5×10^7^). In blue, *E. coli* MG1655 wild type. In red, a strain with a deletion of the *fhuA* gene (DR42). In green, *E. coli* carrying an empty pXG1 plasmid, used as a control. In purple, *E. coli* carrying plasmid pXG1 containing the *llp* gene from phage T5. **Right.** Plaque assay using the same strains and bacteriophages T5 and T4. **B.** Growth curves of an *E. coli* derivative strain DR41 carrying a P*_opvAB_*::*llp*-*gfp* fusion in the presence or absence of T5 phage for 10 hours. Inlet, the same data are represented as a function of time. **C.** GFP fluorescence distribution in DR41 (P*_opvAB_::llp-gfp)* before (t = 0) and after growth in LB containing T5, unknown *E. coli* phages (Ec_Unk_PO1, Ec_Unk_PO2, Ec_Unk_PO3 isolated on *E. coli* MG1655, and Ec_Unk_EM1 isolated on DR28), or colicin M (3 µg/ml, t = 8 h). Data are represented by a dot plot (side scatter versus fluorescence intensity [ON subpopulation size]). All data were collected for 100,000 events per sample.

## Discussion

The challenge of detecting and accurately counting bacteriophages has been around for some time as more and more researchers are interested in characterizing bacteriophages for ecological or biotechnological purposes. Beside the gold standard plaque assay method, a variety of techniques have been developed, each with pros and cons (52, 53). These techniques are based on methodologies such as quantitative PCR (ddPCR or regular qPCR), FISH, electron microscopy, and flow cytometry, and can be classified into three types: culture-dependent, sequence-based, and particle-based (53). Another way to classify these techniques is to consider the characteristics of the enumeration and the additional infection parameters that can be obtained (46). For example, the classical plaque assay can discriminate infectious versus non-infectious particles, gives information on the adsorption rate, is doable without any particular equipment, does not need genome sequencing, but it is difficult to perform in a high-throughput manner. In contrast, a ddPCR-based method allows high throughput analysis but requires knowledge of the phage genome sequence and highly specialized equipment. The information provided is also different as there is no possibility to discriminate infectious versus non-infectious particles while it can discern between different phages in a single assay (52). In the spectra of current methods, flow cytometry-based approaches are among the most sensitive and most published methods. They rely on labelling of the phage genome with a fluorescent dye (54–56), although some label-free protocols have been developed recently (57).

In the present work, we combine culture-based methods with flow cytometry, which allows to quantify with high sensitivity infectious particles only (Fig. 4). Starting with the original biosensor construct in *S. enterica* (Figs. 1, 2) able to detect purified or unpurified phages present in raw water samples (Fig. 3), we expanded our collection of biosensors to detect coliphages. The phase variation *opvAB* locus is present *Salmonella enterica* but neither in *Salmonella bongori* nor in other enteric bacteria (32). In a MG1655 derivative with a complete (reconstructed) LPS, a constitutively expressed *opvAB* operon integrated at the *lac* locus shortens the *E. coli* O-antigen (Fig. 6A). As a consequence, the engineered *E. coli* strain is able to detect LPS-binding phages in a way similar to the *S. enterica* strain (Figs. 2, 6). Of note, the enrichment of the OpvAB^ON^ subpopulation by killing of the OFF subpopulation was not as effective with coliphages as for phages active against *S. enterica*. Nevertheless, as the detection by flow cytometry is very sensitive and reproducible, the enrichment levels achieved were sufficient for a proper detection. The sensitivity was however decreased, perhaps indicating that the coliphages used in the study required an additional component of the outer membrane as receptor, rendering LPS binding less crucial for successful infection. This hypothesis is strengthened by the fact that in the case of the FhuA-binding phages detected using the P_opvAB_-*llp*::*gfp* biosensor the enrichment of the ON subpopulation was much more efficient (Fig. 7). Indeed, it is known that T5 relies essentially on FhuA for a productive lytic cycle, whereas the polymannose decoration of the O-antigen is used as a primary receptor important for primary binding but dispensable for the overall cycle (Fig. 7) (58, 59).

Whole-cell biosensors have been developed to detect a wide range of compounds toxic to bacteria such as antibiotics, heavy metals and toxic chemicals such as arsenic, toluene and phenol (60–64). In all cases, the cell must survive in order to transcend the signal making it quantifiable, and not all bacterial strains are adapted to such detection (65). Indeed, traditional whole-cell sensors have problems detecting compounds with a high toxicity or other external agents that kill the bacterial biosensor cells such as toxins or bacteriophages. In contrast, we show that a biosensor based on a phase-variation regulated system such as the *opvAB* operon is able to detect toxins (Colicin M) and bacteriophages, because the subpopulation which does not confer resistance (OFF) is killed while the subpopulation that confers resistance (ON) survives and becomes enriched, thus allowing the detection of a reporter gene. Furthermore, lysis of the OFF subpopulation increases the amount of phage, speeding up the enrichment of the ON subpopulation and increasing the detection sensitivity. An important add-on value of the method described here is its capacity to discriminate bacteriophages infecting the same host but using different types of receptors (Figs. 6, 7). This property is relevant in the frame of phage therapy when isolation of phages using different receptors is a must to overcome resistance due to mutations in the receptor (66). This work, allowing a rapid discrimination between phages using LPS or FhuA as receptors, could thus be systematically used prior to the assembly of phage cocktails. Moreover, this method could be implemented for other receptors used by bacteriophages as long as a specific inhibitor is available (like Llp for FhuA-binding phage: Fig. 7). For T-even coliphages, for example, a trick might be to use the outer-membrane protein TraT, encoded by the F plasmid, that masks or modifies the conformation of outer-membrane protein A (OmpA), which is the receptor for many T-even-like phages (23, 67, 68). If an inhibitor-encoding gene does not exist for a dedicated receptor, an alternative would be to use the receptor gene itself fused to the *gfp* and controlled by the *opvAB* promoter. A potential drawback of this strategy is that the biosensor might be less sensitive for phage enumeration as the presence of phage would be detected as a decrease in fluorescence. Nevertheless, this alternative procedure to determine the host receptor remains open.

## Acknowledgments

This study was supported by grant BIO2016-75235-P from the Ministerio de Ciencia, Innovación y Universidades of Spain to J.C. and grant ANR-16-CE12-0029-01 from the Agence Nationale de la Recherche to M.A. We are grateful to Elena Puerta-Fernández for advice and discussions, to Rocío Fernández for her help in sample processing, and to Modesto Carballo, Laura Navarro and Cristina Reyes from the Servicio de Biología, CITIUS, Universidad de Sevilla, for help with experiments performed at the facility. We are also very thankful to Artemis Kosta and Hugo Le Guenno at the imaging facility of the Mediterranean Institute of Microbiology (IMM) for TEM imaging, and Denis Duché, LISM Marseille, for the kind gift of colicin M. M.A. is grateful to Nicolas Ginet for critical reading of the manuscript and to the whole phage group @LCB for constant support regarding the phage project at the University of Sevilla.

## References

1. Wommack KE, Colwell RR. 2000. Virioplankton: viruses in aquatic ecosystems. Microbiol Mol Biol Rev MMBR 64:69–114.

2. Roux S, Hallam SJ, Woyke T, Sullivan MB. 2015. Viral dark matter and virus-host interactions resolved from publicly available microbial genomes. eLife 4.

3. Beller L, Matthijnssens J. 2019. What is (not) known about the dynamics of the human gut virome in health and disease. Curr Opin Virol 37:52–57.

4. Lawrence D, Baldridge MT, Handley SA. 2019. Phages and Human Health: More Than Idle Hitchhikers. Viruses 11:587

5. Jover LF, Effler TC, Buchan A, Wilhelm SW, Weitz JS. 2014. The elemental composition of virus particles: implications for marine biogeochemical cycles. Nat Rev Microbiol 12:519– 528.

6. Abeles SR, Pride DT. 2014. Molecular Bases and Role of Viruses in the Human Microbiome. J Mol Biol 426:3892–3906

7. Allen HK, Abedon ST. 2014. Virus ecology and disturbances: impact of environmental disruption on the viruses of microorganisms. Front Microbiol 5:700.

8. Hurwitz BL, Westveld AH, Brum JR, Sullivan MB. 2014. Modeling ecological drivers in marine viral communities using comparative metagenomics and network analyses. Proc Natl Acad Sci U S A 111:10714–10719.

9. Rinke C, Schwientek P, Sczyrba A, Ivanova NN, Anderson IJ, Cheng J-F, Darling A, Malfatti S, Swan BK, Gies EA, Dodsworth JA, Hedlund BP, Tsiamis G, Sievert SM, Liu W-T, Eisen JA, Hallam SJ, Kyrpides NC, Stepanauskas R, Rubin EM, Hugenholtz P, Woyke T. 2013. Insights into the phylogeny and coding potential of microbial dark matter. Nature 499:431–437.

10. Hatfull GF. 2015. Dark Matter of the Biosphere: the Amazing World of Bacteriophage Diversity. J Virol 89:8107–8110.

11. Salmond GPC, Fineran PC. 2015. A century of the phage: past, present and future. Nat Rev Microbiol 13:777–786.

12. Rohwer F, Segall AM. 2015. In retrospect: A century of phage lessons. Nature 528:46– 48.

13. Ofir G, Sorek R. 2018. Contemporary Phage Biology: From Classic Models to New Insights. Cell 172:1260–1270.

14. Peitzman SJ. 1969. Felix d’Herelle and bacteriophage therapy. Trans Stud Coll Physicians Phila 37:115–123.

15. Chanishvili N. 2016. Bacteriophages as Therapeutic and Prophylactic Means: Summary of the Soviet and Post Soviet Experiences. Curr Drug Deliv 13:309–323.

16. Abedon ST, García P, Mullany P, Aminov R. 2017. Editorial: Phage Therapy: Past, Present and Future. Front Microbiol 8:981.

17. Rohde C, Resch G, Pirnay J-P, Blasdel BG, Debarbieux L, Gelman D, Górski A, Hazan R, Huys I, Kakabadze E, Łobocka M, Maestri A, Almeida GM de F, Makalatia K, Malik DJ, Mašlaňová I, Merabishvili M, Pantucek R, Rose T, Štveráková D, Van Raemdonck H, Verbeken G, Chanishvili N. 2018. Expert Opinion on Three Phage Therapy Related Topics: Bacterial Phage Resistance, Phage Training and Prophages in Bacterial Production Strains. Viruses 10:178.

18. Djebara S, Maussen C, De Vos D, Merabishvili M, Damanet B, Pang KW, De Leenheer P, Strachinaru I, Soentjens P, Pirnay J-P. 2019. Processing Phage Therapy Requests in a Brussels Military Hospital: Lessons Identified. Viruses 11:265.

19. Farooq U, Yang Q, Ullah MW, Wang S. 2018. Bacterial biosensing: Recent advances in phage-based bioassays and biosensors. Biosens Bioelectron 118:204–216.

20. Franche N, Vinay M, Ansaldi M. 2016. Substrate-independent luminescent phage-based biosensor to specifically detect enteric bacteria such as E. coli. Environ Sci Pollut Res Int 24:42–51.

21. Vinay M, Franche N, Grégori G, Fantino J-R, Pouillot F, Ansaldi M. 2015. Phage-Based Fluorescent Biosensor Prototypes to Specifically Detect Enteric Bacteria Such as E. coli and Salmonella enterica Typhimurium. PLoS ONE 10:e0131466.

22. Rakhuba DV, Kolomiets EI, Dey ES, Novik GI. 2010. Bacteriophage receptors, mechanisms of phage adsorption and penetration into host cell. Pol J Microbiol Pol Tow Mikrobiol Pol Soc Microbiol 59:145–155.

23. Bertozzi Silva J, Storms Z, Sauvageau D. 2016. Host receptors for bacteriophage adsorption. FEMS Microbiol Lett 363:fnw002.

24. Letarov AV, Kulikov EE. 2017. Adsorption of Bacteriophages on Bacterial Cells. Biochem Biokhimiia 82:1632–1658.

25. Andres D, Hanke C, Baxa U, Seul A, Barbirz S, Seckler R. 2010. Tailspike interactions with lipopolysaccharide effect DNA ejection from phage P22 particles in vitro. J Biol Chem 285:36768–36775.

26. Hu B, Margolin W, Molineux IJ, Liu J. 2015. Structural remodeling of bacteriophage T4 and host membranes during infection initiation. Proc Natl Acad Sci U S A 112:E4919–4928.

27. Arnaud C-A, Effantin G, Vivès C, Engilberge S, Bacia M, Boulanger P, Girard E, Schoehn G, Breyton C. 2017. Bacteriophage T5 tail tube structure suggests a trigger mechanism for Siphoviridae DNA ejection. Nat Commun 8:1953.

28. Wang C, Tu J, Liu J, Molineux IJ. 2019. Structural dynamics of bacteriophage P22 infection initiation revealed by cryo-electron tomography. Nat Microbiol 4:1049.

29. Davies MR, Broadbent SE, Harris SR, Thomson NR, van der Woude MW. 2013. Horizontally Acquired Glycosyltransferase Operons Drive Salmonellae Lipopolysaccharide Diversity. PLoS Genet 9:e1003568.

30. Cota I, Sánchez-Romero MA, Hernández SB, Pucciarelli MG, García-del Portillo F, Casadesús J. 2015. Epigenetic Control of Salmonella enterica O-Antigen Chain Length: A Tradeoff between Virulence and Bacteriophage Resistance. PLoS Genet 11:e1005667.

31. Wahl A, Battesti A, Ansaldi M. 2019. Prophages in Salmonella enterica: a driving force in reshaping the genome and physiology of their bacterial host? Mol Microbiol 111:303–316.

32. Cota I, Blanc-Potard AB, Casadesús J. 2012. STM2209-STM2208 (opvAB): a phase variation locus of Salmonella enterica involved in control of O-antigen chain length. PloS One 7:e36863.

33. Walter M, Fiedler C, Grassl R, Biebl M, Rachel R, Hermo-Parrado XL, Llamas-Saiz AL, Seckler R, Miller S, van Raaij MJ. 2008. Structure of the receptor-binding protein of bacteriophage det7: a podoviral tail spike in a myovirus. J Virol 82:2265–2273.

34. Casjens SR, Leavitt JC, Hatfull GF, Hendrix RW. 2014. Genome Sequence of Salmonella Phage 9NA. Genome Announc 2:e00531–14.

35. Datsenko KA, Wanner BL. 2000. One-step inactivation of chromosomal genes in Escherichia coli K-12 using PCR products. Proc Natl Acad Sci USA 97:6640–6645.

36. Blattner FR, Plunkett G, Bloch CA, Perna NT, Burland V, Riley M, Collado-Vides J, Glasner JD, Rode CK, Mayhew GF, Gregor J, Davis NW, Kirkpatrick HA, Goeden MA, Rose DJ, Mau B, Shao Y. 1997. The complete genome sequence of Escherichia coli K-12. Science 277:1453–1474.

37. Stevenson G, Neal B, Liu D, Hobbs M, Packer NH, Batley M, Redmond JW, Lindquist L, Reeves P. 1994. Structure of the O antigen of Escherichia coli K-12 and the sequence of its rfb gene cluster. J Bacteriol 176:4144–4156.

38. Browning DF, Wells TJ, França FLS, Morris FC, Sevastsyanovich YR, Bryant JA, Johnson MD, Lund PA, Cunningham AF, Hobman JL, May RC, Webber MA, Henderson IR. 2013. Laboratory adapted Escherichia coli K-12 becomes a pathogen of Caenorhabditis elegans upon restoration of O antigen biosynthesis. Mol Microbiol 87:939–950.

39. Hautefort I, Proença MJ, Hinton JCD. 2003. Single-copy green fluorescent protein gene fusions allow accurate measurement of Salmonella gene expression in vitro and during infection of mammalian cells. Appl Environ Microbiol 69:7480–7491.

40. Olivenza DR, Nicoloff H, Antonia Sánchez-Romero M, Cota I, Andersson DI, Casadesús J. 2019. A portable epigenetic switch for bistable gene expression in bacteria. Sci Rep 9:11261.

41. Wilkinson RG, Gemski P, Stocker BA. 1972. Non-smooth mutants of Salmonella typhimurium: differentiation by phage sensitivity and genetic mapping. J Gen Microbiol 70:527–554.

42. Smith HO, Levine M. 1964. TWO SEQUENTIAL REPRESSIONS OF DNA SYNTHESIS IN THE ESTABLISHMENT OF LYSOGENY BY PHAGE P22 AND ITS MUTANTS. Proc Natl Acad Sci U S A 52:356–363.

43. Chan RK, Botstein D, Watanabe T, Ogata Y. 1972. Specialized transduction of tetracycline resistance by phage P22 in Salmonella typhimurium. II. Properties of a high-frequency-transducing lysate. Virology 50:883–898.

44. Torreblanca J, Casadesús J. 1996. DNA adenine methylase mutants of Salmonella typhimurium and a novel dam-regulated locus. Genetics 144:15–26.

45. Buendía-Clavería AM, Moussaid A, Ollero FJ, Vinardell JM, Torres A, Moreno J, Gil-Serrano AM, Rodríguez-Carvajal MA, Tejero-Mateo P, Peart JL, Brewin NJ, Ruiz-Sainz JE. 2003. A purL mutant of Sinorhizobium fredii HH103 is symbiotically defective and altered in its lipopolysaccharide. Microbiol Read Engl 149:1807–1818.

46. Ackermann H-W. 2009. Basic phage electron microscopy. Methods Mol Biol Clifton NJ 501:113–126.

47. Flayhan A, Wien F, Paternostre M, Boulanger P, Breyton C. 2012. New insights into pb5, the receptor binding protein of bacteriophage T5, and its interaction with its Escherichia coli receptor FhuA. Biochimie 94:1982–1989.

48. Braun V, Killmann H, Herrmann C. 1994. Inactivation of FhuA at the cell surface of Escherichia coli K-12 by a phage T5 lipoprotein at the periplasmic face of the outer membrane. J Bacteriol 176:4710–4717.

49. Pedruzzi I, Rosenbusch JP, Locher KP. 1998. Inactivation in vitro of the Escherichia coli outer membrane protein FhuA by a phage T5-encoded lipoprotein. FEMS Microbiol Lett 168:119–125.

50. Decker K, Krauel V, Meesmann A, Heller KJ. 1994. Lytic conversion of Escherichia coli by bacteriophage T5: blocking of the FhuA receptor protein by a lipoprotein expressed early during infection. Mol Microbiol 12:321–332.

51. Washizaki A, Yonesaki T, Otsuka Y. 2016. Characterization of the interactions between Escherichia coli receptors, LPS and OmpC, and bacteriophage T4 long tail fibers. Microbiology Open 5:1003–1015.

52. Morella NM, Yang SC, Hernandez CA, Koskella B. 2018. Rapid quantification of bacteriophages and their bacterial hosts in vitro and in vivo using droplet digital PCR. J Virol Methods 259:18–24.

53. Baran N, Goldin S, Maidanik I, Lindell D. 2018. Quantification of diverse virus populations in the environment using the polony method. Nat Microbiol 3:62–72.

54. Brussaard CPD. 2004. Optimization of procedures for counting viruses by flow cytometry. Appl Environ Microbiol 70:1506–1513.

55. Carreira C, Staal M, Middelboe M, Brussaard CPD. 2015. Counting viruses and bacteria in photosynthetic microbial mats. Appl Environ Microbiol 81:2149–2155.

56. de la Cruz Peña MJ, Martinez-Hernandez F, Garcia-Heredia I, Lluesma Gomez M, Fornas Ò, Martinez-Garcia M. 2018. Deciphering the Human Virome with Single-Virus Genomics and Metagenomics. Viruses 10:113.

57. Ma L, Zhu S, Tian Y, Zhang W, Wang S, Chen C, Wu L, Yan X. 2016. Label-Free Analysis of Single Viruses with a Resolution Comparable to That of Electron Microscopy and the Throughput of Flow Cytometry. Angew Chem Int Ed Engl 55:10239–10243.

58. Braun V, Wolff H. 1973. Characterization of the receptor protein for phage T5 and colicin M in the outer membrane of E. coli B. FEBS Lett 34:77–80.

59. Heller K, Braun V. 1982. Polymannose O-antigens of Escherichia coli, the binding sites for the reversible adsorption of bacteriophage T5+ via the L-shaped tail fibers. J Virol 41:222–227.

60. Ivask A, Rõlova T, Kahru A. 2009. A suite of recombinant luminescent bacterial strains for the quantification of bioavailable heavy metals and toxicity testing. BMC Biotechnol 9:41.

61. Tauriainen S, Karp M, Chang W, Virta M. 1998. Luminescent bacterial sensor for cadmium and lead. Biosens Bioelectron 13:931–938.

62. Prévéral S, Brutesco C, Descamps ECT, Escoffier C, Pignol D, Ginet N, Garcia D. 2016. A bioluminescent arsenite biosensor designed for inline water analyzer. Environ Sci Pollut Res Int 24:25–32.

63. Shingler V, Moore T. 1994. Sensing of aromatic compounds by the DmpR transcriptional activator of phenol-catabolizing Pseudomonas sp. strain CF600. J Bacteriol 176:1555–1560.

64. Tecon R, Beggah S, Czechowska K, Sentchilo V, Chronopoulou P-M, McGenity TJ, van der Meer JR. 2010. Development of a multistrain bacterial bioreporter platform for the monitoring of hydrocarbon contaminants in marine environments. Environ Sci Technol 44:1049–1055.

65. Brutesco C, Prévéral S, Escoffier C, Descamps ECT, Prudent E, Cayron J, Dumas L, Ricquebourg M, Adryanczyk-Perrier G, de Groot A, Garcia D, Rodrigue A, Pignol D, Ginet N. 2016. Bacterial host and reporter gene optimization for genetically encoded whole cell biosensors. Environ Sci Pollut Res Int 24:52–65.

66. Bai J, Jeon B, Ryu S. 2019. Effective inhibition of Salmonella Typhimurium in fresh produce by a phage cocktail targeting multiple host receptors. Food Microbiol 77:52–60.

67. Riede I, Eschbach ML. 1986. Evidence that TraT interacts with OmpA of Escherichia coli. FEBS Lett 205:241–245.

68. Labrie SJ, Samson JE, Moineau S. 2010. Bacteriophage resistance mechanisms. Nat Rev Microbiol 8:317–327.

69. Manoil C, Beckwith J. 1985. TnphoA: a transposon probe for protein export signals. Proc Natl Acad Sci U S A 82:8129–8133.

70. Simon R, Priefer U, Pühler A. 1983. A Broad Host Range Mobilization System for In Vivo Genetic Engineering: Transposon Mutagenesis in Gram Negative Bacteria. Bio/Technology 1:784–791.

71. Edwards RA, Keller LH, Schifferli DM. 1998. Improved allelic exchange vectors and their use to analyze 987P fimbria gene expression. Gene 207:149–157.

72. Urban JH, Vogel J. 2007. Translational control and target recognition by Escherichia coli small RNAs in vivo. Nucleic Acids Res 35:1018–1037.

